# How multilevel competition drives cooperation in stigmergic systems

**DOI:** 10.64898/2026.10.28.762141

**Authors:** Thomas Bassanetti, Ramón Escobedo, Stéphane Cezera, Maxime Delacroix, Diego Álvarez, Miguel Moscoso, Adrien Blanchet, Clément Sire, Guy Theraulaz

## Abstract

Cooperation through stigmergy relies on reliable information left in a shared environment, yet individuals may benefit from misleading others. How competition within and between groups shapes this conflict remains unclear. We investigated this question using the *Stigmer Game*, in which human participants interact exclusively through ratings that modify a shared informational environment, under seven payment rules. Participants were classified as collaborators, neutrals or defectors. Between-group competition produced the highest collaboration, whereas within-group competition favoured defection. Adding between-group competition partly restored cooperation despite continued rivalry within groups, although this effect depended on reward distribution. A quantitative analysis showed that within-group and rank-based competitive pressures best described differences in behavioural composition across payment rules. A genetic algorithm further revealed distinct stationary strategy compositions under each payment rule that were largely independent of initial conditions and robust to perturbations. These results reproduced the opposing effects of between-group and within-group competition observed experimentally, while high-lighting differences between immediate behavioural responses and the strategy distributions favoured by each rule’s incentive structure. Defectors remained present under the mutation regime, illustrating the vulnerability of shared information to exploitation. Our findings show how the organization of competitive incentives shapes cooperation and information reliability in decentralized collective systems.

## 1 Introduction

Cooperation is a defining feature of biological and human societies. From microbial biofilms and insect colonies to human organizations, individuals frequently contribute to collective outcomes while pursuing their own interests. Understanding how cooperation emerges and persists despite conflicts of interest is therefore a central challenge in evolutionary biology, behavioural ecology, economics and complex systems science [1, 2]. Classical explanations emphasize kin selection, reciprocal altruism, indirect reciprocity and multilevel selection as mechanisms aligning individual and collective interests [3–7]. Complementary approaches highlight self-organization, through which local interactions generate coordinated collective behaviour without centralized control [2, 8–10]. Together, these perspectives connect evolutionary cooperation with collective intelligence, whereby groups solve problems through distributed information processing [11–13]. A prominent mechanism of decentralized coordination is stigmergy, originally introduced to explain nest construction in social insects [14, 15]. Individuals interact indirectly by modifying their shared environment, leaving traces that influence subsequent behaviour. Pheromone trails in ants illustrate how such interactions generate feedback loops and collective organization [16, 17]. Stigmergic coordination extends to pedestrian trails, collaborative human activities, online communities and distributed artificial systems [2, 18–20]. Across these systems, local environmental information supports the emergence of collective patterns without requiring global knowledge or centralized control [21, 22].

Although stigmergy explains how decentralized coordination emerges, it does not explain why individuals should contribute reliable information when their interests conflict. Individuals may exploit information generated by others while contributing little or deliberately misleading their partners [23]. Such behaviour can undermine collective information processing and compromise the benefits of stigmergic coordination. Maintaining cooperation therefore requires understanding how individual incentives influence the production and reliability of shared information [11]. Previous studies have demonstrated that reward structures affect cooperation and defection, but have primarily examined interactions within individual groups. This leaves open the question of how cooperation changes when groups are embedded in larger competitive environments [6, 24, 25]. Addressing this question requires considering selective pressures acting at different organizational levels [26, 27].

Experimental approaches provide a means of investigating how incentive structures affect cooperation. Most studies rely on public goods games, prisoner’s dilemma games or common-pool resource paradigms [28, 29]. Although these approaches have advanced our understanding of social dilemmas, they primarily examine cooperation through the production or sharing of resources. They rarely address cooperation mediated by environmental information [30]. Yet such information is central to many collective systems. Ant colonies coordinate foraging through pheromone trails [8, 17], while ants and termites organise construction by modifying their environment [31–33]. Human communities similarly generate ratings, digital traces and shared databases that guide collective decisions [34, 35]. In these systems, cooperation involves producing reliable information that benefits other individuals rather than directly transferring resources [36]. Understanding how incentives affect the reliability of this shared information is therefore essential to explaining stigmergic cooperation.

The *Stigmer Game* provides a controlled experimental framework for investigating this problem [37]. Participants explore a spatial environment containing cells with different values and assign ratings to the cells they visit. These ratings become visible to other group members as coloured traces, progressively forming a shared environmental memory that guides subsequent exploration. Participants can collaborate by assigning ratings that accurately reflect cell values or defect by providing misleading information that disadvantages their partners. Previous experiments demonstrated that payment rules strongly influence the emergence of collaborative, neutral and defective behavioural profiles [37]. Equal rewards promoted extensive cooperation, whereas competition based on individual ranking increased deceptive behaviour. These findings established that changing incentives alone can reshape the information collectively generated through stigmergic interactions.

However, previous *Stigmer Game* experiments considered only situations without explicit competition or with competition restricted to members of the same group. Biological and human collectives frequently experience competition at several organizational levels simultaneously [38]. Individuals compete with their groupmates for resources, status or rewards, while groups compete with one another for collective benefits. Multilevel evolutionary game theory predicts that these pressures have opposing effects on cooperation [39, 40]. Within-group competition favours individual advantage and can encourage defection, whereas betweengroup competition rewards collective performance and can promote cooperation among group members [24, 25, 41]. Importantly, these pressures may interact nonlinearly, producing outcomes that cannot be predicted from either level of competition considered separately [6, 13]. Whether these theoretical predictions extend to cooperation mediated by shared environmental information remains largely unexplored.

Here, we investigate how multilevel competition shapes stigmergic cooperation using the *Stigmer Game*. Seven payment rules manipulate competition within groups, between groups or at both levels. We quantify their effects on human behaviour by measuring the proportions of collaborators, neutrals and defectors. We then use a genetic algorithm, in which experimentally characterized behavioural profiles serve as fixed agent strategies, to examine strategy distributions favoured by each payment rule. Combining these approaches allows us to distinguish immediate behavioural responses from the strategy distributions obtained with the genetic algorithm and investigate how competing incentives influence cooperation. Our findings show that cooperation depends on the balance between competitive pressures operating at different organisational levels and on the distribution of individual rewards [42]. By connecting stigmergic coordination with multilevel selection theory, this study provides a framework for understanding how incentive structures affect the reliability of shared information in biological collectives, human organizations and distributed artificial systems [11, 21, 22].

## 2 Materials and Methods

### 2.1 Experimental design

Experiments were conducted at the TSE Experimental Laboratory using the *Stigmer Game*, a computer-based collective search task designed to investigate cooperation through indirect information sharing [37]. The game was implemented through a custom-built interactive web application developed partly in collaboration with Andil (https://www.andil.fr). Before each session, participants provided informed consent and received instructions explaining the task, payment conditions and guarantees of anonymity. They were also asked to switch off their mobile phones. Participants were seated in individual cubicles, assigned at random and identified by anonymous codes, which prevented direct interactions between them; see figure 1(*b,c*).

**Figure 1:**
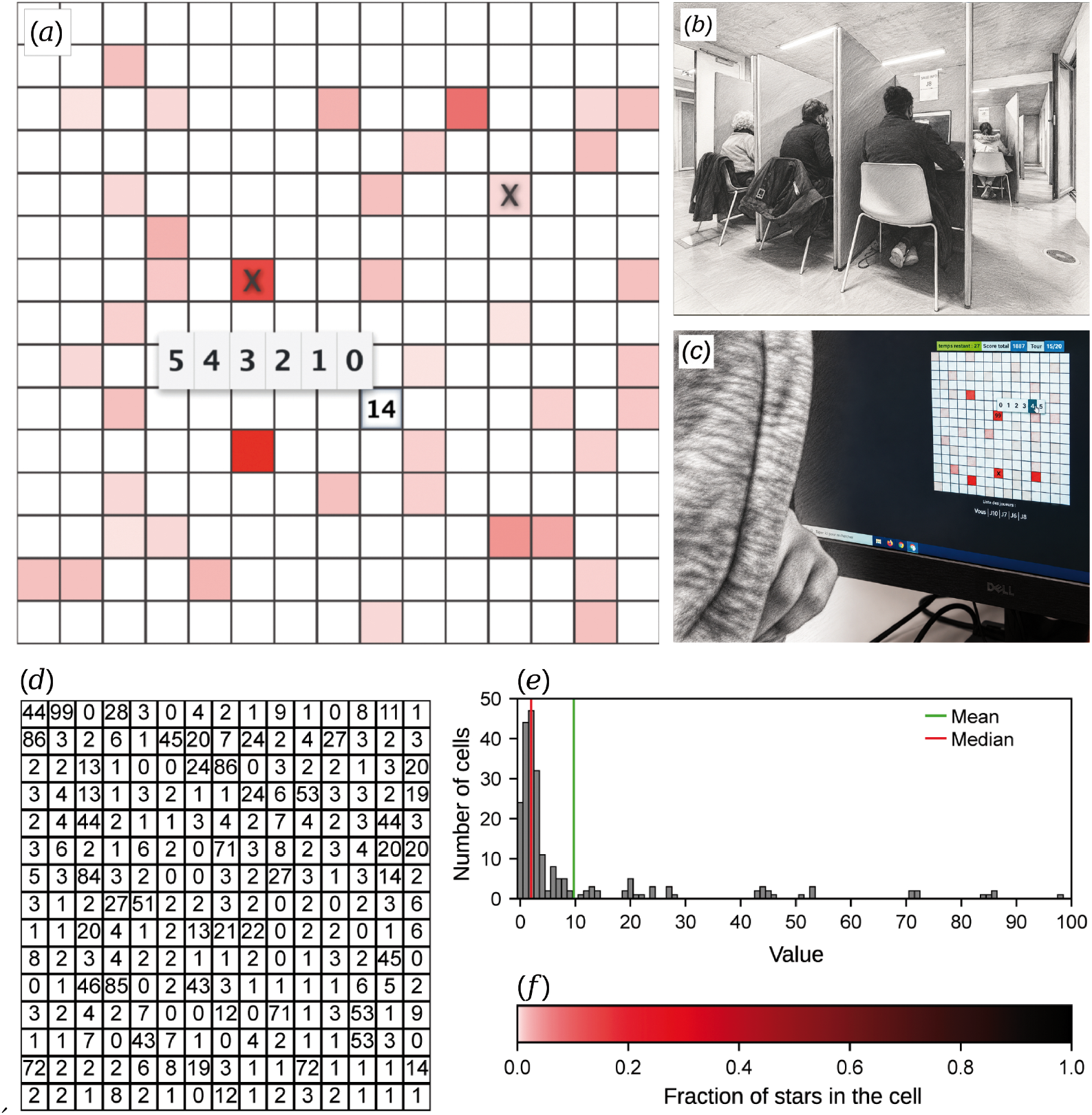
Experimental setup and visual representation of shared information. (*a*) Screenshot of the game interface at round *t* = 10. The participant has already visited and rated two cells, indicated by black crosses, and has just selected a third cell with a value of *V* = 14. The participant must assign a rating from zero to five stars, while the score increases by 14 points. (*b*) Experimental room showing the individual cubicles used to prevent direct interactions between participants. (*c*) Computer interface used during the experiments. (*d*) Example of a 15 × 15 grid used in the experiments. (*e*) Distribution of the 225 integer cell values *V*, ranging from 0 to 99, shared by all grids but randomly rearranged between games. (*f* ) Colour scale representing the fraction of stars accumulated by each cell relative to the total number of stars assigned across the grid. White indicates either an unvisited cell or a cell that has received only zero-star ratings. Increasingly darker shades of red indicate larger fractions of accumulated stars, with black corresponding to the maximum fraction. Colour intensity follows a square-root transformation to improve visual contrast.

Each participant explored a 15 × 15 grid of 225 cells displayed on their computer screen, with each cell associated with a hidden value *V* ranging from 0 to 99. All grids contained the same set of 225 values, randomly rearranged between games. Participants in both groups explored the same grid of cell values, but each group did so independently, with no information shared between groups. Examples of the grids were provided during the instructions, and figure 1(*d,e*) shows a representative grid and the distribution of cell values.

At each round, participants selected a cell, observed its value and assigned it a rating from zero to five stars (figure 1(*a*)). The ratings contributed to a shared visual representation of the grid, allowing participants to exchange information indirectly. Rather than seeing the individual ratings assigned by other group members, participants observed the colour of each cell, which reflected the fraction of stars it had received relative to the total number of stars assigned across the grid. Colours ranged from white, indicating no accumulated stars, through progressively darker shades of red to black, with colour intensity scaled to the square root of this fraction to improve visual contrast (figure 1(*f* )). All cells were initially white, and cells that had never been visited remained visually indistinguishable from those that had received only zero-star ratings. As participants explored and rated cells, these evolving colour patterns formed a shared environmental memory that could guide subsequent exploration, reproducing the principle of stigmergic coordination [37].

Each experimental session involved 10 participants and lasted approximately one hour. Participants were randomly divided into two groups of five, with group membership remaining fixed throughout the session. Each individual started by playing two games of twenty rounds alone, during which the only available information came from their own activity. These preliminary games familiarised participants with the task and provided a baseline measure of their behaviour in the absence of social information [37]. Each group then played approximately 10 collective games of 20 rounds, with a new random arrangement of cell values in each game.

During collective games, participants could use the shared colour patterns generated by members of their own group but had no access to information produced by the other group. Direct communication was not permitted. Participants accumulated points according to the values of the cells they visited and received feedback on their individual performance. Their final monetary reward was determined by one of seven payment rules, with the same rule applied throughout each session. Between seven and ten independent sessions were conducted for each payment rule, using different participants in each session.

### 2.2 Experimental conditions

The seven payment rules determine whether competition operates within groups, between groups, at both levels, or is absent. When the payment rule involves between-group competition, the scores of both groups are used jointly to determine the individual rewards. The task itself and the information available to players remained unchanged across rules.

Under **NC-Equal** (*No Competition, Equal reward*), each player receives 10 €, independently of score, rank or group performance. This condition provides a baseline in which monetary rewards generate neither withingroup nor between-group competition.

Under **NC-Score** (*No Competition, reward by Score*), payment is proportional to the individual score. Participants were informed that a score corresponding to the average performance measured in previous sessions would yield a payment of approximately 11 €. Because one player’s performance does not reduce another player’s reward, this condition contains no explicit competition for a limited reward.

Under **WG** (*Within-Group competition only*), players compete exclusively with members of their own group. They are ranked according to their individual score within the group. The first-ranked player receives 20 €, the second 15 €, and the remaining three players 10 € each. The performance of the other group has no effect on these rewards.

Under **BG** (*Between-Group competition only*), the two groups compete according to the mean individual score of their five members. Each member of the group with the highest mean score receives 20 €, whereas each member of the other group receives 10 €. Rewards therefore depend exclusively on collective performance and are identical for all members of the same group. Individual rank within a group has no direct effect on payment.

Three **BW** conditions (*Between- and Within-Group competition*) combine both levels of competition. Participants from both groups are ranked together according to their individual score. Under **BW-Step**, the five highest-ranked players receive 15 € each and the remaining five receive 8 € each. Under **BW-Linear**, payments decrease linearly with rank, according to the formula 16 ™ rank, from 15 € for the highest-ranked player to 6 € for the lowest-ranked player. Under **BW-Podium**, the first-ranked player receives 20 €, the second and third receive 15 €, and the remaining seven players receive 10 € each.

Mean payments per player are approximately 10, 11, 13, 15, 11.5, 10.5 and 12 € for `NC-Equal`, `NC-Score`, `WG`, `BG`, `BW-Step`, `BW-Linear` and `BW-Podium`, respectively. Under the `NC-Score` rule, payments were normalized within each session to keep the total payment for the 10 participants fixed at 110 €. The reward received by participant *i* was therefore calculated as

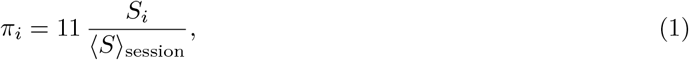

where *S*_*i*_ is the participant’s score and ⟨*S*⟩_session_ is the mean score of the 10 participants in that session.

Consequently, the final payment was not determined solely by the participant’s own score but also depended weakly on the performance of the other participants in the session. Participants were not informed of this normalization and therefore had no information that could induce strategic competition with other players. From the participants’ perspective, `NC-Score` thus remained a non-competitive condition in which increasing another player’s score did not appear to reduce their own expected reward.

### 2.3 Behavioural profiles classification

Players are classified according to how the ratings they assign depend on the true values of the visited cells, following the behavioural classification previously introduced in [37]. For each player, the mean rating assigned to cells of value *V* is approximated by a linear function *u*_0_ + *u*_1_ × 5*V/*99. Players with *u*_1_ ≥ 0.5 are *collaborators*, because their ratings convey reliable information about cell quality. Players with *u*_1_ *<* ™0.5 are *defectors*, because their ratings convey misleading information, and players with intermediate slopes are *neutrals*. The two thresholds correspond to the minima of the empirical distribution of *u*_1_.

Behavioural composition under each payment rule *r* is quantified by the fractions of collaborators *C*_*r*_, neutrals *N*_*r*_ and defectors *D*_*r*_, with *C*_*r*_ + *N*_*r*_ + *D*_*r*_ = 1.

### 2.4 Competitive pressure indicators

The incentive structures of the seven payment rules can be compared through three measures of competitive pressure, *C*_within_, *C*_between_, and *C*_rank_, that quantify reward differentiation within groups, between groups, and across the full ranking of all players, respectively:

- Within-group competitive pressure is quantified as the mean dispersion of rewards within the two groups,

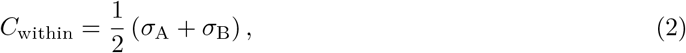

where *σ*_A_ and *σ*_B_ are the standard deviations of individual rewards within groups A and B. Thus, *C*_within_ = 0 when all members of each group receive the same reward and increases as reward differences among group members become larger.
- Between-group competitive pressure is quantified as the difference between the mean rewards received by the winning and losing groups,

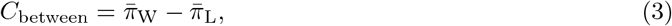

where 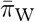 and 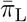 denote the mean rewards of the winning and losing groups, respectively. This indicator is zero when group membership has no effect on mean reward and increases as the reward difference between groups becomes larger.
- Rank-based competitive pressure across all players is quantified as

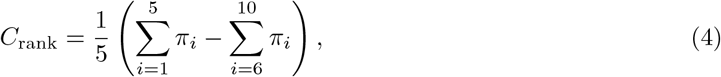

where *π*_*i*_ is the reward received by the *i*-th player when the 10 are ranked according to their individual score.

For payment rules with predetermined rewards, *C*_within_ is calculated directly from the reward structure. When *C*_between_ or *C*_rank_ depend on how individual ranks are distributed between groups, their expected values are estimated using a null model in which ranks are randomly assigned to the two groups and averaged over 10^5^ realizations. Under `NC-Score`, rewards depend on the scores actually obtained and could not be specified in advance. The three indicators are therefore calculated from the mean rewards observed at each rank across experimental sessions. These mean values are, in decreasing order, 13.38, 13.13, 12.63, 11.75, 11.13, 10.88, 10.63, 9.88, 9.50, and 8 €.

The three indicators are then used in regression models to determine which combination best accounts for the observed behavioural profiles. Indicators are first normalized by the mean payment of each rule 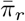, and then standardized across the seven rules. The standardized indicator is defined as

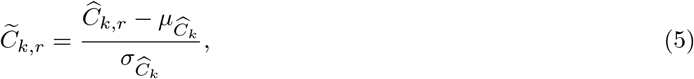

for *k* ∈ {within, between, rank}, where 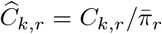 is the normalized indicator, and 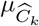 and 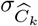 are the mean and standard deviation of 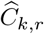 across payment rules.

We fit the seven possible linear models predicting *C*_*r*_ and *D*_*r*_ from all possible combinations of the three standardized indicators: three single-indicator models, three two-indicator models, and one model with all three indicators. Model fit is evaluated by the normalized prediction error

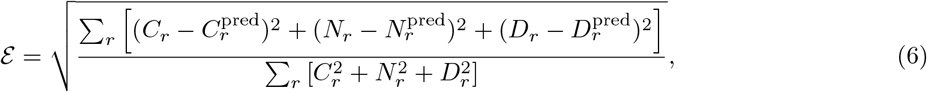

where the sum is over the seven payment rules. All indicators are standardized before fitting, so regression coefficients can be compared directly across indicators. The complete comparison among candidate models, together with fitted coefficients and prediction errors, is provided in the electronic supplementary material.

### 2.5 Genetic algorithm

To identify the distribution of behavioural strategies favoured by each payment rule, we used a genetic algorithm adapted from [43], with a population of *N* = 100 agents and a mutation rate of *µ* = 0.1.

Each agent carries one of three fixed behavioural strategies, collaborator, neutral, or defector, derived from the experimental profiles identified in [37]. Agent behaviour remains unchanged until death and replacement, and is governed by a stochastic agent-based model comprising two components: a visit strategy determining which cells are selected, and a rating strategy determining how visited cells are rated according to their value. In the genetic algorithm, all agents have the same visit strategy and differ only by their rating strategy: collaborators generate informative ratings, defectors generate systematically misleading ratings, and neutrals assign ratings independently of cell value (figure S1). The complete behavioural model, including the functional forms and parameters governing cell exploration, revisiting and rating, is described in detail in the electronic supplementary material.

The algorithm runs for *N*_gen_ = 1000 generations, each consisting of two phases, an interaction phase and a reproduction phase. During the interaction phase, each agent plays the *Stigmer Game* a large number of times (*N*_e_ = 1000). Agent fitness is updated after each game according to the payment rule, via an exponential memory rule with parameter *ε* = 0.1. In the reproduction phase, all agents survive except a small fraction *p*_death_ = 0.05 of low-fitness agents, which are replaced by copies of high-fitness agents selected from the upper 40% of the population. Each offspring mutates with probability *µ* = 0.1 to one of the three strategies according to a given mutation spectrum. Two spectra are considered: a random spectrum (*µ*_C_, *µ*_D_) = (1*/*3, 1*/*3), and an experimental spectrum (*µ*_C_, *µ*_D_) = (0.84, 0.03) derived from the behavioural baseline observed under `NC-Equal`.

Each run is continued until the distribution of fractions converged to a stationary stable state. Stationary fractions are averaged over the last *N*_av_ = 50 generations and over *N*_samp_ = 50 independent replicates. Convergence is confirmed by initializing from multiple population compositions, including homogeneous single-strategy populations and random mixtures. Stability is assessed by replacing at least 60% of the stationary population with agents carrying another strategy and determining whether the population returned to the same stationary composition.

The robustness of the rule ordering with respect to the fraction of collaborators is verified by systematic parameter sweeps: *µ* in [0, 0.35], *ε* in [0, 1], and *p*_death_ in [0.02, 0.2]. A complete description of the algorithm, parameter choices, and convergence analyses is provided in the electronic supplementary material.

## 3 Results

### 3.1 Multilevel competition reshapes cooperation

We first examine how the seven payment rules affected individual performance and the information that participants contributed to their group. Performance is quantified using the normalized score *S* and the probability 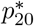 of having found one of the highest-value cells at round 20, the last of a game. Behavioural profiles are classified independently of individual performance.

Individual performance differed markedly across payment rules (figure 2). `NC-Equal` produced the lowest mean normalized score, with ⟨*S*⟩ = 0.24, compared with values close to or above 0.4 under the other six rules. This difference reflects the absence of any incentive to maximise individual performance under `NC-Equal`. Participants instead explored the grid extensively, as previously observed in [37]. Consistent with this exploratory behaviour, 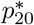 reached 0.35 under `NC-Equal`, compared with values below 0.2 under the other rules. Notably, `NC-Equal` was the only condition in which this probability exceeded both the random and sequential exploration benchmarks (see Section A of the electronic supplementary material for their derivation).

**Figure 2:**
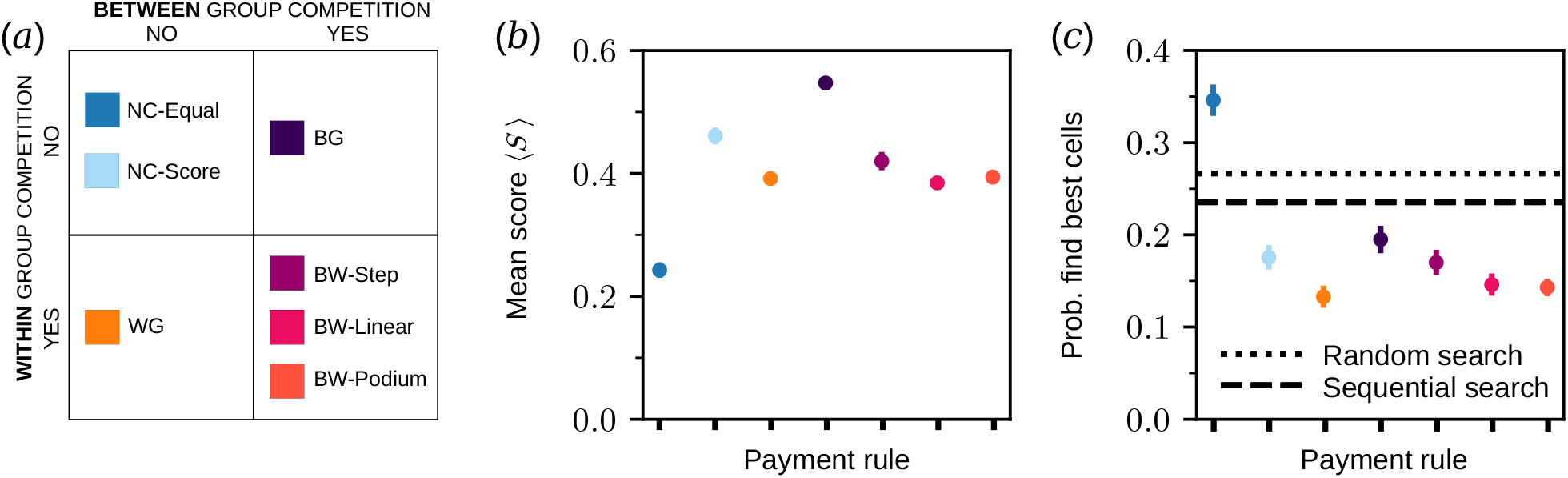
Individual performance across the seven payment rules. (*a*) Competition structure across the seven payment rules. The seven colour-coded payment rules are positioned according to the presence or absence of within-group and between-group competition, distinguishing non-competitive (NC-Equal, NC-Score), within-group (WG), between-group (BG), and combined competition (BW-Step, BW-Linear, BW-Podium) conditions. (*b*) Mean normalized score ⟨*S*⟩. (*c*) Probability of having found one of the nine highest-value cells at the final round (*t* = 20). The black dashed and dotted lines indicate the expected probabilities under random and sequential exploration, respectively.

Among the conditions that rewarded performance, `BG` and `NC-Score` yielded the highest scores. Neither rule imposed explicit competition among members of the same group. Under `BG`, however, individual rewards depended entirely on collective success, whereas `NC-Score` rewarded participants according to their own performance. Conditions containing explicit within-group competition generally produced lower performance. This pattern was strongest under `WG` and remained apparent under the three BW conditions. Thus, rewarding performance alone did not determine how efficiently participants used the shared information. Performance also depended on whether participants’ interests were aligned with or opposed to those of their groupmates.

The clearest effect of the reward structure emerged from the behavioural profiles (table 1). The three conditions without explicit within-group competition contained the largest fractions of collaborators. Under `NC-Equal`, 84% were collaborators, 13% were neutrals, and only 3% were defectors. Cooperation became even stronger under `BG`, where collaborators represented 93% of participants and no defectors were observed. Competition between groups therefore did not simply preserve cooperation within groups. When collective performance determined individual rewards, it produced the highest level of collaboration observed across all experimental conditions.

**Table 1:**
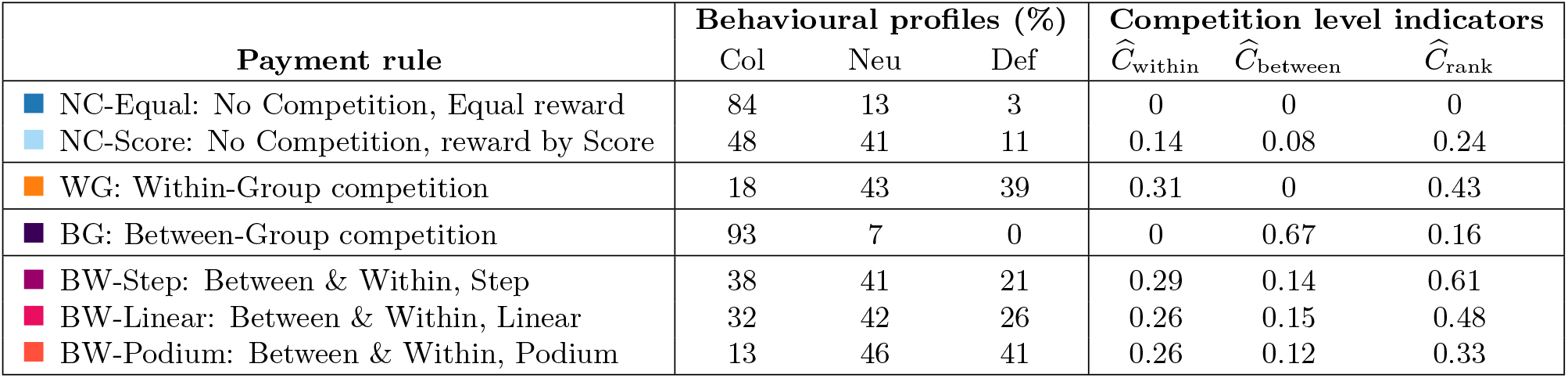
Behavioural profiles and competitive pressures across payment rules. Percentages of collaborators (Col), neutrals (Neu), and defectors (Def) observed under each experimental condition, together with the normalized competitive pressure measures *Ĉ*_within_, *Ĉ*_between_, and *Ĉ*_rank_, as defined in Section 2.

| **Payment rule** | **Behavioural profiles (%)** |  |  | **Competition level indicators** |  |  |
|---|---|---|---|---|---|---|
|  | Col | Neu | Def | Ĉ<sub>within</sub> | Ĉ<sub>between</sub> | Ĉ<sub>rank</sub> |
| NC-Equal: No Competition, Equal reward | 84 | 13 | 3 | 0 | 0 | 0 |
| NC-Score: No Competition, reward by Score | 48 | 41 | 11 | 0.14 | 0.08 | 0.24 |
| WG: Within-Group competition | 18 | 43 | 39 | 0.31 | 0 | 0.43 |
| BG: Between-Group competition | 93 | 7 | 0 | 0 | 0.67 | 0.16 |
| BW-Step: Between & Within, Step | 38 | 41 | 21 | 0.29 | 0.14 | 0.61 |
| BW-Linear: Between & Within, Linear | 32 | 42 | 26 | 0.26 | 0.15 | 0.48 |
| BW-Podium: Between & Within, Podium | 13 | 46 | 41 | 0.26 | 0.12 | 0.33 |

The pattern reversed when participants competed against members of their own group. Under `WG`, collaborators represented only 18% of participants, whereas defectors increased to 39%, revealing a sharp collapse of cooperation when within-group competition acted alone. Importantly, introducing between-group competition alongside within-group competition partially restored cooperation. Collaborators increased to 38% under `BW-Step` and 32% under `BW-Linear`, while defectors decreased to 21% and 26%, respectively. This protective effect was not universal, however, as `BW-Podium` produced only 13% collaborators and 41% defectors. The fraction of neutrals remained comparatively stable across `WG` and the three BW conditions, ranging from 41% to 46%. Thus, within-group competition strongly shifts behaviour from collaboration towards defection, but competition between groups can counteract this effect when the reward structure sufficiently aligns individual success with collective performance.

`NC-Equal` and `NC-Score` provide informative exceptions. Under `NC-Equal`, 84% of participants collaborate despite the complete absence of monetary incentive to do so. Under `NC-Score`, only 48% collaborate, despite the absence of explicit competition. These fractions differ strongly from those observed under `NC-Equal`. Because rewards under `NC-Score` depended only on individual performance, participants could behave competitively even though improving another player’s score did not directly reduce their own reward. These two distinctions becomes important below, because the experimentally observed response to `NC-Score` cannot be fully represented by a model in which selection depends only on monetary payoff.

### 3.2 Competitive pressures predict behavioural responses

The large behavioural differences among payment rules raise a simpler question. Which properties of the reward structure actually determine whether participants collaborate or defect? We addressed this question by characterizing each payment rule with three normalized measures of competitive pressure. *C*_within_ quantifies reward differentiation within groups, *C*_between_ quantifies reward differentiation between groups, and *C*_rank_ quantifies reward differentiation associated with individual ranking across all participants. These measures depend only on the incentive structure and not on the behavioural classification itself, thereby providing a compact description of the competitive pressures generated by each payment rule.

The experimental data reveal a strong relationship between competitive pressures and behaviour (table 1). The fraction of collaborators decreases strongly as *C*_within_ increases. This effect is clearest under `WG`, where within-group competition acts without an opposing between-group incentive and the fraction of collaborators falls to 18%. However, the BW conditions show that the effect of within-group competition depends on the broader competitive context. When between-group competition is introduced simultaneously, cooperation can be partially restored, reaching 38% under `BW-Step` and 32% under `BW-Linear`. Thus, *C*_within_ represents the main pressure opposing stigmergic cooperation, whereas competition between groups can counterbalance this pressure by realigning individual interests with collective performance.

To quantify these relationships, we compared the seven possible linear models constructed from all combinations of the standardized measures 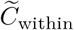, 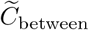 and 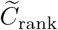. These comprised three single-predictor models, three two-predictor models and one model containing all three predictors. The models predicted the fractions of collaborators and defectors, while the fraction of neutrals followed from *N*_*r*_ = 1 ™ *C*_*r*_ ™ *D*_*r*_. Among the single-predictor models, *C*_within_ provided the best prediction of the observed behavioural profiles, with an error *ε* = 0.18 (figure 3a). It could not, however, distinguish `NC-Equal` from `BG`, because both have *C*_within_ = 0.

**Figure 3:**
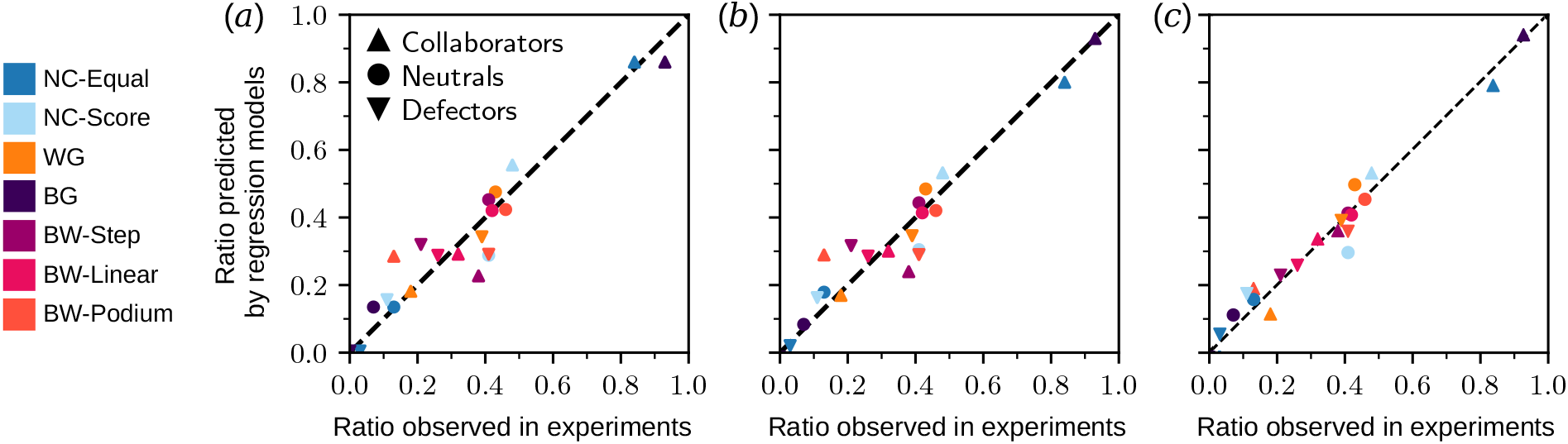
Regression model predictions compared with experimental behavioural profiles. Predicted fractions of collaborators, neutrals, and defectors plotted against the corresponding fractions observed experimentally for each payment rule. Predictions are shown for (*a*) the within-group competitive pressure model 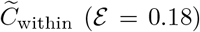, (*b*) the model combining within- and between-group competitive pressures 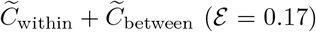, and (*c*) the model combining within-group and rank-based competitive pressures 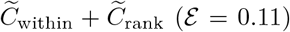. The dashed diagonal indicates perfect agreement between predicted and observed behavioural fractions.

Adding *C*_between_ to *C*_within_ produced only a marginal improvement, reducing the prediction error to *ε* = 0.17 (figure 3b). Its main contribution was to distinguish `BG` from `NC-Equal`. The coefficients associated with *C*_between_ were typically 10 to 20 times smaller than those associated with *C*_within_. Moreover, when all three competitive pressures were included in the model, the coefficients associated with *C*_between_ remained small and changed sign. These results indicate that *C*_between_ carries little independent explanatory power in the linear regression, despite the strong behavioural effect observed when between-group competition is introduced experimentally.

The best prediction was obtained by combining *C*_within_ and *C*_rank_, which reduced the prediction error to *ε* = 0.11 (figure 3c). Expressed relative to the mean behavioural composition observed across payment rules, the fitted model was

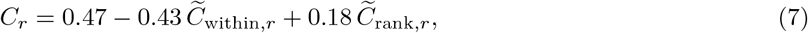

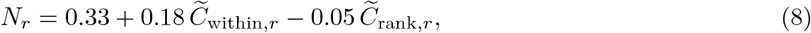

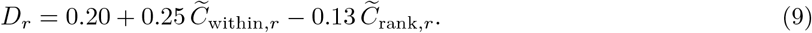

Because both predictors were standardized, their coefficients can be compared directly. The coefficients associated with *C*_within_ were approximately two to three times larger in absolute value than those associated with *C*_rank_, confirming that within-group competitive pressure is the dominant predictor of behavioural composition. Increasing *C*_within_ strongly reduced the fraction of collaborators while increasing both neutrals and defectors. This effect was stronger for defectors than for neutrals, indicating that increasing withingroup competitive pressure shifts behaviour primarily towards information manipulation rather than towards neutral information sharing. By contrast, increasing *C*_rank_ increased collaboration and reduced defection. Its effect on neutrals was weak, with a coefficient of only ™0.05.

The contribution of *C*_rank_ is particularly informative. Although weaker than *C*_within_, its inclusion substantially improved predictions for intermediate behavioural fractions between 20% and 40%. It also provided a more useful description of the global competitive structure than *C*_between_. The latter is dominated by the particularly high value associated with `BG` and therefore mainly separates this condition from `NC-Equal`, whereas *C*_rank_ discriminates competitive pressure across the full set of payment rules.

These results show that the seven incentive structures need not be considered as unrelated experimental treatments. Most of their behavioural effects can be summarized by a low-dimensional description of competitive pressure. *C*_within_ provides the dominant contribution and strongly opposes stigmergic cooperation, whereas *C*_rank_ accounts for additional variation in behavioural composition. Importantly, the weak independent contribution of *C*_between_ in the regression does not imply that between-group competition has little behavioural effect. The experimental comparison between `WG` and the BW conditions shows that introducing between-group competition can partially restore cooperation when within-group competition is already present. The regression instead indicates that this effect is better captured across the seven payment rules by the global reward structure summarized by *C*_rank_. Together, these results identify a low-dimensional incentive architecture that predicts immediate behavioural responses. We next used a genetic algorithm to determine the strategy distributions favoured by each payment rule from its incentive structure alone.

### 3.3 Ranking of payment rules by their ability to promote cooperation

To assess how the seven payment rules favour different behavioural strategies, we used a genetic algorithm to compare the strategy distributions obtained under each rule. This approach provides a ranking based on the incentives created by the payment rules, independently of the experimental observations. The genetic algorithm incorporates a stochastic behavioural model of agents playing the *Stigmer Game*, adapted from [37] and described in detail in Section C of the electronic supplementary material.

The algorithm combines three processes that determine the distribution of behavioural strategies. Genetic drift introduces random changes in strategy frequencies, while mutation continually introduces strategies according to a specified mutation spectrum. Selection favours strategies according to the fitness differences generated by each payment rule. Together, these processes determine the stationary strategy distributions obtained by the algorithm, allowing us to compare the extent to which different payment rules favour cooperation.

Figure 4 shows the changes in strategy frequencies and the stationary distributions obtained with the genetic algorithm under each payment rule, using a uniform mutation spectrum. Under the two NC rules, collaborator frequencies remain relatively close to those expected from the mutation spectrum, although the stationary population under `NC-Score` is dominated by neutrals. In contrast, `BG` strongly favours collaboration, with collaborators accounting for approximately 86% of the stationary population, whereas `WG` strongly favours defection, with defectors reaching 89%. The three BW rules produce intermediate levels of collaboration, with defectors outnumbering collaborators in all three cases. The fraction of collaborators decreases from `BW-Step` to `BW-Linear` and then to `BW-Podium`. The dynamics under `NC-Score` are particularly noteworthy. Collaborators initially increase in frequency while defectors are still present, before gradually giving way to a stationary population dominated by neutrals. The mechanisms underlying this transient increase in collaboration and the subsequent dominance of neutrals are discussed in Section 4. Simulations using a biased mutation spectrum yield the same qualitative ordering of payment rules (figure S2), indicating that the differences in their ability to sustain collaboration are primarily driven by the selective pressures associated with each rule rather than by the mutation spectrum alone.

**Figure 4:**
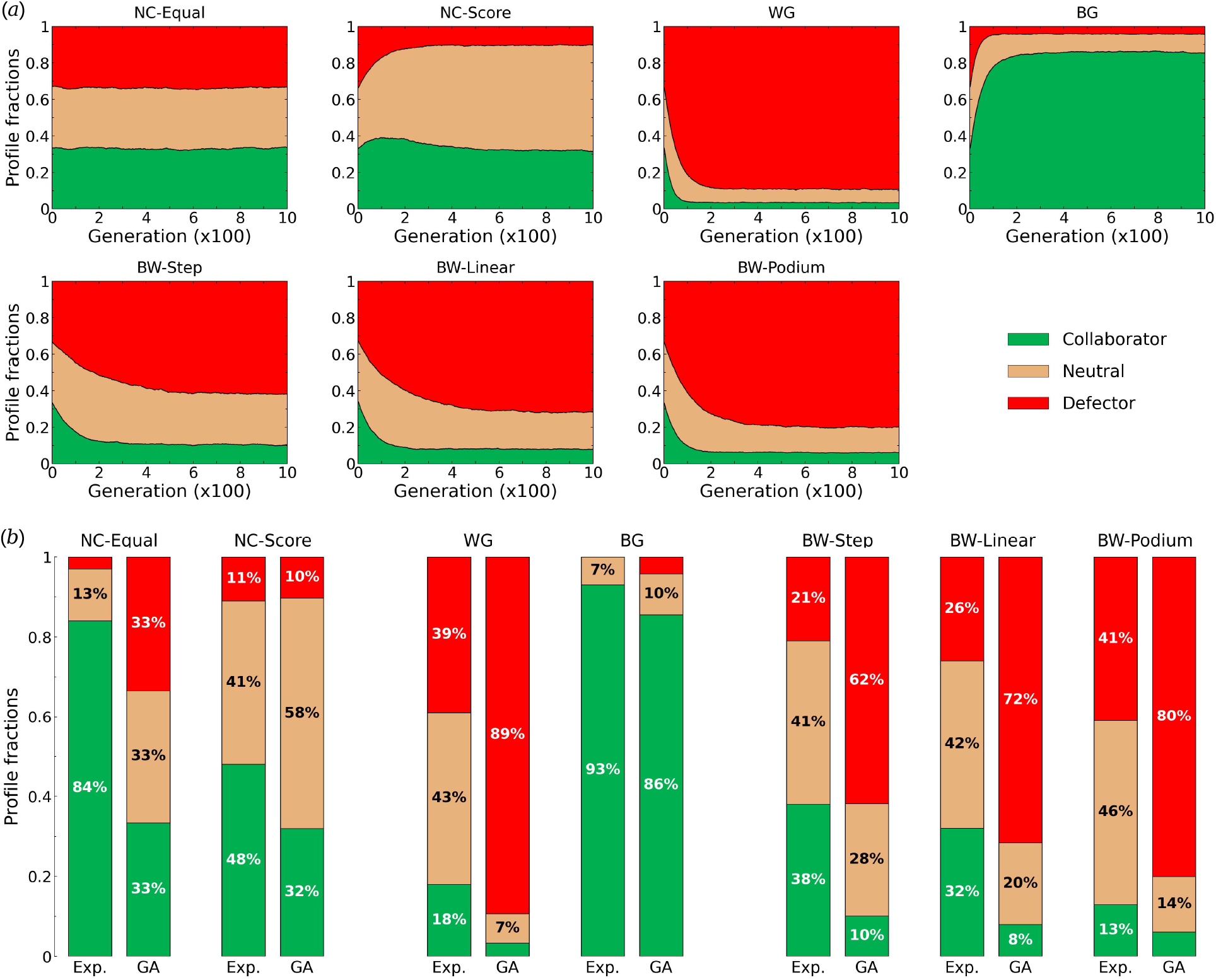
Genetic algorithm dynamics and comparison of stationary strategy distributions with experimental behavioural profiles under the seven payment rules, using a uniform mutation spectrum. (*a*) Changes in strategy frequencies over successive generations of the genetic algorithm, starting from random initial strategy compositions. (*b*) Comparison of the asymptotic fractions of collaborators, neutrals and defectors obtained with the genetic algorithm (GA) with the corresponding fractions measured experimentally (Exp.) for each of the seven payment rules. Green areas show the fraction of collaborators *C*_*r*_, red areas show 1 ™ *D*_*r*_, where *D*_*r*_ is the fraction of defectors, brown area between both shows the fraction of neutrals 1 ™ *C*_*r*_ ™ *D*_*r*_. Parameter values of the GA: *N* = 1000, *µ* = 0.1, *N*_gen_ = 1000, *ε* = 0.1, *p*_death_ = 0.05, *p*_parents_ = 0.40. Individuals are replaced using stochastic universal sampling (SUS), with mutation probabilities 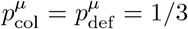. Average of *N*_samp_ = 500 replicas, final state averaged over the last *N*_av_ = 100 generations. See figure S2 in the electronic supplementary material for biased mutations, figure S5 for different configurations of the initial population, and figure S6 for the invasion tests.

Varying the mutation rate *µ* reveals how the strategy distributions obtained by the algorithm change as mutations become increasingly rare (figure S3). As *µ* → 0, populations tend towards fixation of a single behavioural strategy, with the strategy favoured depending on the payment rule. Under `BG`, collaborators approach fixation, whereas defectors dominate under `WG` and the three BW rules. Under `NC-Score`, the population tends towards fixation of neutrals, while `NC-Equal` leads to stochastic fixation of any of the three strategies, consistent with the absence of selective differences between them. Thus, in the absence of a continuous supply of collaborators through mutation, none of the rules except `BG` consistently favours the persistence of cooperation. Despite these differences in the strategies favoured at low mutation rates, the relative ordering of the payment rules in terms of collaborator frequency remains unchanged over the investigated range *µ* ∈ [0, 0.35]. Strategy frequencies vary gradually with *µ*, with no apparent discontinuities or changes in the relative ordering of the payment rules.

The relative ordering of the seven payment rules is also preserved when varying the fitness memory parameter *ε* ∈ [0, 1] and the population renewal rate *p*_death_ ∈ [0.02, 0.2] (figure S4), indicating that the differences in their ability to promote cooperation are robust across the parameter ranges investigated.

The stationary strategy distributions are also largely insensitive to initial population composition and resilient to substantial perturbations. Simulations starting from widely different strategy frequencies, including homogeneous populations and random mixtures, converge towards similar stationary distributions under each payment rule (figure S5). Furthermore, invasion experiments in which 60% of the population is replaced by agents adopting a different strategy show that populations recover their characteristic stationary compositions following the perturbation (figure S6).

Taken together, these results show that the seven payment rules produce distinct and robust stationary strategy distributions. Their relative ability to sustain cooperation is primarily determined by the fitness differences associated with each rule and remains largely unchanged across the algorithm parameters and initial conditions examined.

## 4 Discussion

Stigmergy enables individuals to coordinate their activities indirectly by modifying a shared environment. However, the effectiveness of this form of collective coordination depends on the reliability of the information left by others. When individual interests conflict with collective interests, individuals may benefit from providing misleading information, potentially undermining cooperation. Understanding how competition within and between groups shapes these conflicting incentives is therefore essential to explaining how cooperation emerges and persists in stigmergic systems. We investigated this question using the *Stigmer Game* under seven payment rules that independently vary the intensity and scope of competition, combining behavioural experiments, quantitative regression models, and a genetic algorithm that identifies the strategy distributions favoured by each rule from its incentive structure alone. Together, these approaches show that between-group competition strongly promotes cooperation, whereas within-group competition favours defection. However, the outcome also depends on how rewards are distributed among individuals. Competition therefore shapes not only individual behaviour but also the reliability of the shared information on which collective intelligence depends.

The most striking result is the contrast between within-group and between-group competition. In the experiments, `BG` yielded 93% collaborators and no defectors, whereas under `WG`, collaboration fell to 18% and defection rose to 39%. The genetic algorithm suggests that this pattern reflects the structure of the payment rules, with collaborators approaching fixation under `BG` and defectors under `WG` as the mutation rate approaches zero. These findings are consistent with multilevel selection theory, which highlights the opposing selective pressures operating within and between groups [6, 24, 27, 44], and extend earlier work using the *Stigmer Game* [37, 45]. Crucially, between-group competition did more than reward collective performance. By making individual rewards depend on group success, it created incentives to contribute useful information to the shared environment. In contrast, within-group competition favoured strategies that exploited this information without making equally useful contributions, undermining the collective benefits of stigmergic coordination.

Introducing between-group competition can partially restore cooperation even when individuals continue to compete within their own group. In our experiments, the fraction of collaborators increased from 18% under `WG` to 38% under `BW-Step` and 32% under `BW-Linear`, while defection decreased accordingly. These results are consistent with a central prediction of multilevel selection theory, namely that competition between groups can favour cooperation despite the opposing selective pressures operating within groups [6, 24, 27].

The genetic algorithm provides further insight into how the distribution of rewards shapes this balance between competition levels. It consistently predicts the ordering `BW-Step` *>* `BW-Linear` *>* `BW-Podium` in the fraction of collaborators, suggesting that the accessibility of higher rewards to the different behavioural strategies plays an important role. Under `BW-Step`, the five highest-ranked participants across both groups receive the higher payment, allowing collaborators, who tend to rank below defectors, a greater opportunity to benefit than under `BW-Podium`, where the largest rewards are concentrated among the top three ranks. `BW-Linear` falls between these two conditions, with a gradual decrease in payment as rank declines. Because mutation continually reintroduces all three behavioural strategies, their stationary frequencies reflect the balance between this influx and the selective advantages created by each reward structure. The differences in collaboration between the three BW conditions thus suggest that the extent to which collaborators can access higher-ranked rewards helps determine how effectively between-group competition counteracts selection for defection within groups.

However, the introduction of between-group competition did not always favour cooperation. In the experiments, `BW-Podium` yielded only 13% collaborators and 41% defectors, a lower level of cooperation than under `WG`. Concentrating the highest rewards among a small number of top-ranked individuals may intensify competition and encourage deceptive behaviour, even when individuals also compete as members of a group. By contrast, the genetic algorithm consistently placed all three BW rules between the strongly cooperation-promoting `BG` condition and the defector-dominated `WG` condition. These findings suggest that between-group competition can offset the effects of within-group rivalry, but that its ability to do so depends critically on how rewards are allocated.

The differences are small and may reflect the fact that human behaviour is not driven by monetary incentives alone. Social preferences, perceptions of fairness and sensitivity to relative performance may influence how individuals respond to different reward structures. A useful extension of the model would therefore be to examine how individual and collective incentives jointly shape behaviour, for example by introducing a fitness function of the form *απ*_individual_ + *βπ*_collective_, where *α* and *β* control the relative contributions of individual and collective rewards.

Our quantitative analysis further shows that the way competitive incentives are structured matters more than simply whether competition occurs within or between groups. Within-group competitive pressure alone accounted for much of the variation in experimental behaviour (*ε* = 0.18), and including between-group pressure brought only a marginal improvement (*ε* = 0.17). The best fit was obtained by combining within-group and rank-based competitive pressures (*ε* = 0.11). This does not mean that between-group competition has little influence, as illustrated by the high level of cooperation observed under `BG`. Rather, rank-based pressure appears to capture more effectively how differences in reward allocation shape behaviour across the seven payment rules. These results highlight the importance of how individual incentives are aligned with collective interests in determining the benefits of cooperation [1, 11].

The `NC-Score` condition reveals a further distinction between the formal structure of incentives and how participants respond to them. Although this rule involved no explicitly stated competition for a limited reward, only 48% of participants behaved as collaborators, compared with 84% under `NC-Equal`. Linking monetary rewards to individual performance may therefore be sufficient to encourage more self-interested behaviour, even in the absence of explicit competition between participants. Whether this response reflects a perceived sense of competition or other motivational factors remains to be investigated.

The genetic algorithm explains this apparent paradox. Although `NC-Score` does not explicitly reward relative performance, it nevertheless generates selection between behavioural strategies within the algorithm. Monetary rewards depend on individual scores rather than rankings, but the genetic algorithm selects individuals for reproduction and replacement according to their relative fitness. As a result, differences in individual performance translate into selective advantages even under this nominally non-competitive rule. The resulting changes in strategy frequencies are non-monotonic. When defectors are abundant, collaborators initially gain an advantage because their reliable ratings help counteract the misleading information introduced by defectors, allowing collaboration to increase in frequency. As defectors become scarce, however, neutrals acquire a small but persistent advantage. Their behaviour combines elements of both defection and collabo-ration. By leaving some high-value cells weakly marked, they retain information that remains less accessible to other group members, while repeatedly visiting and rating other cells allows them to contribute to the shared information available to the group. This balance between withholding and sharing information can make neutrality more advantageous than consistent collaboration.

Although modest, this advantage is measurable. In a group initially composed of four collaborators, replacing one collaborator with a neutral increases the neutral’s mean score by 0.7%, while reducing the mean score of the remaining collaborators by 2.3%. As the mutation rate approaches zero, this small selective advantage favours the fixation of neutrals, making `NC-Score` the only payment rule under which neither collaboration nor defection is consistently favoured in this limit.

Together, the experimental findings and genetic algorithm results suggest that the effects of monetary incentives on cooperation cannot be understood from payment rules alone. They also depend on how behavioural strategies shape the production, sharing and exploitation of information within the group.

Defectors can persist even under conditions that strongly favour cooperation. This may arise partly from an asymmetry in stigmergic information sharing. A defector surrounded by collaborators can benefit from reliable information accumulated in the shared environment while contributing misleading ratings. Conversely, a collaborator surrounded by defectors faces degraded information without receiving comparable benefits from the contributions of others. This asymmetry creates opportunities for defectors to exploit collective information, even in highly cooperative populations [45]. However, their persistence does not necessarily imply a selective advantage under every payment rule. Under `BG`, which most strongly favours collaboration, the genetic algorithm predicts a small but non-zero fraction of defectors maintained by the continuous introduction of new strategies through mutation.

These findings have implications for collective systems that depend on information contributed by their members. Stigmergy enables decentralized coordination through persistent environmental traces [15, 18, 19]. However, these traces become less useful when individuals have incentives to manipulate them. Similar problems arise in online ratings, collaborative databases and other digital environments where shared information guides subsequent decisions [11, 34]. Our results suggest that maintaining reliable information requires more than encouraging participation. Incentive structures must also limit the advantages gained from misleading contributions. Mechanisms that reward information accuracy or penalise deliberate misinformation could complement between-group competition. The *Stigmer Game* offers a controlled framework for testing such interventions before extending them to more complex collective systems.

Several extensions could clarify the mechanisms underlying these results. Providing real-time information about the competing group’s performance would test whether explicit awareness of between-group competition strengthens cooperation, and would introduce strategic plasticity absent from the present framework, bridging the gap between fixed behavioural profiles and reinforcement learning models. Changing group membership between games could reveal how stable social relationships influence behavioural strategies. Varying individual and collective rewards continuously would also help map the transitions between cooperation and defection. Such experiments could establish whether critical thresholds separate distinct behavioural regimes and identify reward structures that best preserve reliable information.

Overall, our study shows that the organization of competition plays a central role in shaping cooperation in stigmergic systems. Between-group competition promotes reliable information sharing, whereas within-group competition generally favours defection. However, their combined effects depend on how individual rewards are distributed and how behavioural strategies interact over time. The genetic algorithm establishes that this ordering is intrinsic to the incentive structure of the rules, independently of the behavioural responses of the participants. The persistence of defectors also highlights a fundamental limitation of stigmergic cooperation, as individuals can exploit shared information while contributing misleading signals. Stigmergy provides a powerful mechanism for decentralized coordination, but it does not guarantee that the information generated collectively will remain reliable [37]. Our findings extend the view of collective intelligence as a public good [11], whose benefits depend on the contributions of individuals and their willingness to share reliable information. They suggest that effective collective intelligence not only requires mechanisms for exchanging and integrating information, but also incentive structures that align individual interests with collective success.

## Ethics

The aims and procedures of the experiments were approved by the Ethics Committee of the Toulouse School of Economics (TSE). All participants provided written consent for their participation.

## Data

The data and all codes that support the findings of this study will be openly available on Figshare.

## Authors’ contributions

The conception and design of the work were carried out by TB, CS, and GT. TB and MD were responsible for the creation of the experimental software used in the work. Data acquisition was performed by TB, SC, RE, AB, CS, and GT. TB, RE, CS, and GT contributed to the data analysis, while data interpretation was conducted by TB, CS, and GT. The genetic algorithm was conceived and designed by RE and CS, and its implementation, analysis, and interpretation were carried out by TB, RE, DA, MM, CS, and GT. The manuscript was drafted by RE and GT. All authors reviewed and approved the final version of the manuscript.

## Electronic Supplementary Material

### A Expected performance of simple search strategies

We provide two theoretical benchmarks for individual search performance: the probability of finding a specific target cell by round *t* by random search or sequential search.

A player visits *N*_turns_ cells per round from a grid containing *N* cells. Under random search, the player selects distinct cells uniformly at random during each round. Cells visited in previous rounds can be visited again. The probability of not finding a given target cell during one round is 1 ™ *N*_turns_*/N* . As searching events independent, the probability of not finding the cell at rount *t* is (1 ™ *N*_turns_*/N* )^*t*^, where the superindex *t* denotes a power, not a time index. Thus,

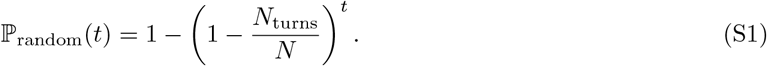

Under sequential search, the player visits only unexplored cells, so the maximum number of rounds is 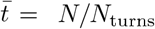. At round 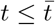, the number of different cells visited is *t* × *N*_turns_, so

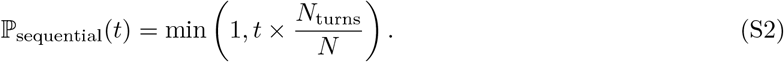

### B Stochastic agent-based model

The behaviour of agents in the genetic algorithm was generated by the stochastic agent-based model previously developed for the *Stigmer Game* [37]. The model contains two components. The visit strategy determines which cells an agent selects, whereas the rating strategy determines how the agent rates a visited cell as a function of its value.

#### B.1 Visit strategy

During the first round (*t* = 1), agents have no prior information and therefore select three cells at random. From the second round onwards (*t >* 1), the three cells visited by an agent are selected through a combination of exploitation and exploration.

For each of the three cells *i* = 1, 2, 3, the agent can revisit the *i*th highest-value cell found during the previous round, whose value is *V*_*i*_(*t* ™ 1). The probability of revisiting this cell is

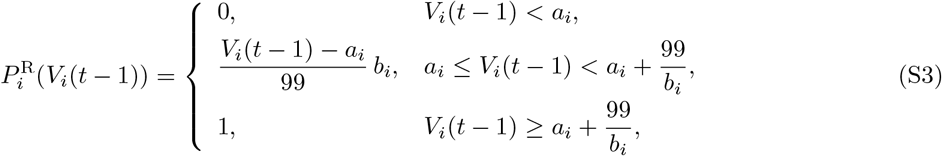

where *a*_*i*_ and *b*_*i*_ *>* 0 are model parameters. An agent therefore never revisits the *i*th best cell when its value is below *a*_*i*_ and always revisits it when its value exceeds *a*_*i*_ + 99*/b*_*i*_, provided that this threshold lies within the possible range of cell values. Between these thresholds, revisit probability increases linearly with cell value.

When the agent does not revisit a previously selected cell, it explores the grid using the social information accumulated in the shared environment. The probability of selecting cell *c* at round *t* is

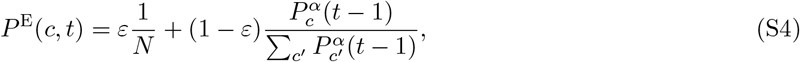

where *P*_*c*_(*t* ™ 1) denotes the cumulative fraction of stars assigned to cell *c* up to round *t* ™ 1. The parameter *ε* ∈ (0, 1) controls the balance between random exploration and the use of stigmergic information, whereas *α >* 0 determines the sensitivity to differences in accumulated ratings. Larger values of *α* increase the preference for highly rated cells.

If the selected cell has already been chosen during the current round or is one of the cells carried over from the preceding round, another cell is drawn. Together, the revisit and exploration rules provide a parsimonious description of the balance between individual exploitation, random exploration and stigmergic information use.

#### B.2 Rating strategy

After visiting a cell of value *V*, an agent assigns a rating *s* ∈ {0, 1, …, 5} according to the probability distribution *P*_*s*_(*V* ). The rating strategy depends only on cell value and is independent of the round, the number of cells already visited during that round and the current stigmergic information associated with the cell (figure S1).

Experimental participants predominantly assigned either 0 or 5 stars, whereas ratings from 1 to 4 stars occurred less frequently and with similar probabilities [37]. We therefore assumed equal probabilities for these four intermediate ratings. For *s* = 1, 2, 3, 4,

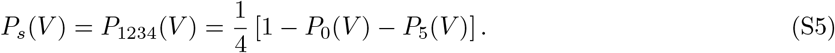

The probabilities of assigning 0 or 5 stars were represented either by sigmoid-like functions,

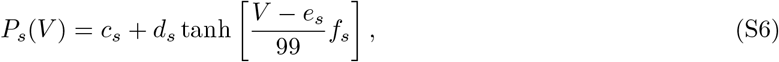

where *c*_*s*_, *d*_*s*_ *>* 0, *e*_*s*_ and *f*_*s*_ are parameters, or by linear functions,

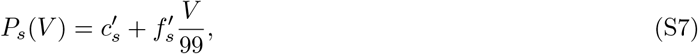

where 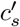 and 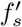 are parameters. These functional forms reproduce the experimentally observed dependence of rating behaviour on cell value while remaining sufficiently flexible to represent different behavioural strategies.

**Figure S1:**
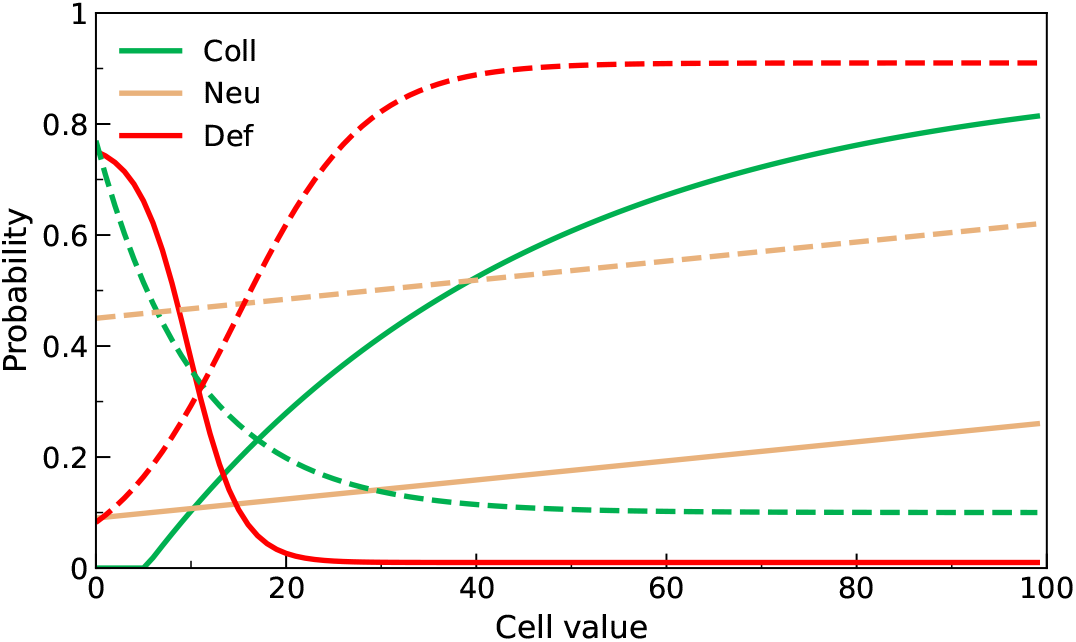
Behavioural profiles of the three strategies. Probability of assigning 5 stars (solid lines) or 0 stars (dashed lines) to a cell as a function of its value *V* ∈ [0, 99], for collaborators (green), neutrals (brown), and defectors (red) [37]. Parameters: collaborator *s* = 0: (*c, d, e, f* ) = (1113.4, 1113.3, ™84.5, ™4.75); *s* = 5: (™1051.9, 1052.8, ™304.5, 1.24). Neutral *s* = 0: (*c*^*′*^, *d*^*′*^) = (0.45, 0.17); *s* = 5: (0.09, 0.17). Defector *s* = 0: (0.45, 0.46, 14.8, 7.34); *s* = 5: (0.39, 0.38, 9.8, ™18.49).

#### B.3 Collaborator, neutral and defector strategies

The three strategies used in the evolutionary simulations were derived from the behavioural profiles identified experimentally [37]. Collaborators preferentially assigned high ratings to high-value cells and low ratings to low-value cells, thereby generating informative stigmergic traces. Defectors showed the opposite relationship and systematically degraded the shared information. Neutral agents displayed little or no systematic relationship between cell value and rating.

These analytical profiles defined the rating probabilities used by each agent type in the genetic algorithm. The visit strategy was governed by the same stochastic framework described above, allowing fitness differences to emerge from repeated interactions between behavioural strategies under the different payment rules.

### C Genetic algorithm

#### C.1 Overview of the genetic algorithm

The genetic algorithm operates on a population of fixed size *N* = 100 over a large number of successive generations.

Each agent adopts one of the three behavioural strategies, collaborator, neutral, or defector, which remains fixed throughout its lifetime. Each generation comprises two successive phases, the interaction phase and the reproduction phase. During the interaction phase, agents repeatedly play the *Stigmer Game*, accumulating fitness according to one of the seven payment rules considered. During the reproduction phase, a fraction of the population is replaced by removing low-fitness agents and introducing new agents which are copies of high-fitness agents. Offspring inherit the strategy of their parent, and eventually undergo a mutation that can result in a change of behavioural strategy.

i. Interaction phase: The interaction phase consists of a series of *N*_e_ = 1000 encounters. Before each encounter, the population is randomly partitioned into *N/*10 = 10 pairs of groups of five players. Then, every group plays a *Stigmer* game of 20 rounds, whose result is associated with the one of the group with which it has been paired. Every agent plays exactly *N*_e_ games per generation. After the *g*-th game, the fitness *f*_*i*_ of every agent *i* is updated according to an exponential memory rule,

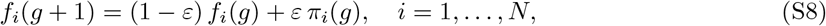

where *π*_*i*_(*g*) is the payment obtained by agent *i* at game *g* and *ε* ∈ (0, 1) is a memory parameter. Fitness values are never normalised or reset during the dynamics. An agent has a weighted history of all its games, where recent games weight more than earlier ones.
ii. Reproduction phase: Following the interaction phase, *N*_death_ = 5 agents are replaced by a copy of a parent drawn from the top *N*_parents_ = 40% of the population ranked by fitness, keeping the population size constant. Agents selected for removal are drawn from the entire population using stochastic universal sampling (SUS), with selection weights proportional to their antifitness, defined as *f*_max_ ™ *f*, where *f*_max_ is the maximum fitness in the population. This procedure makes agents with lower fitness more likely to be removed. For each replacement, a parent is selected uniformly at random, with replacement, from the top *N*_parents_ = 40% of agents ranked by fitness in the population before removal. Each offspring inherits both the behavioural strategy and fitness of its parent. With probability *µ*, a mutation occurs and the offspring is assigned a new strategy sampled according to the mutation probabilities 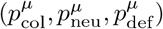. The sampled strategy may coincide with that of the parent, so mutations without a change in strategy are included. When a mutation occurs, the offspring’s fitness is reset to the mean fitness of the population before removal. The mutation probabilities are either random (1*/*3 each profile), or biased based on the behavioural frequencies measured under the `NC-Equal` payment rule: (0.84, 0.13, 0.03). Replacing five agents per generation corresponds to a population renewal rate of *p*_death_ = 5%. This rate is chosen to preserve fitness differences accumulated during the interaction phase, and fast enough to allow the population to reach its stationary state within *N*_gen_ = 1000 generations.

This cycle is repeated for *N*_gen_ = 1000 generations in a population of *N* = 100 agents. After a transient whose duration depends on the selection strength and the replacement and mutation rates, the algorithm reaches a stationary distribution of behavioural strategies.

#### C.2 Fitness update

Let *s*_*j*_ denote the score of agent *j, r*_*j*_ its rank within its group of five, and *R*_*j*_ its overall rank among the ten agents across both groups, with *R*_*j*_ = 1 denoting the player with the highest score. After each game, the fitness of each agent is updated according to equation (S8), with the reward *π*_*j*_ determined by the payment rule as specified below.

**NC-Equal:** Fitness remains unchanged. This rule exerts no selective pressure and serves as a *null baseline*.

**NC-Score:** The reward is proportional to the individual score, normalised by the maximum attainable score *S*_max_ = *N*_*t*_ × (99 + 86 + 86) = 5420, where *N*_*t*_ = 20 is the number of rounds:

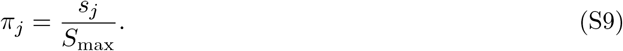

**WG:** The reward is determined by the agent’s rank within its group,

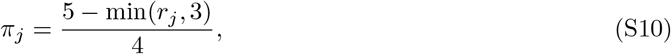

yielding *π* = 1, 0.75, 0.5, 0.5, 0.5 for ranks 1 to 5, respectively.

**BG:** The reward depends exclusively on the relative performance of the two groups, measured by their mean scores:

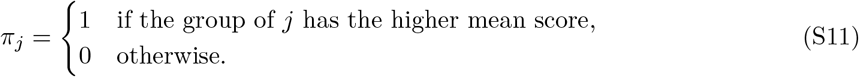

**BW-Step:** The reward is determined by the agent’s overall rank among the ten agents, with a stepwise difference between the top and bottom halves of the ranking:

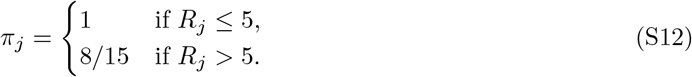

**BW-Linear:** The reward decreases linearly with the agent’s overall rank among the ten agents:

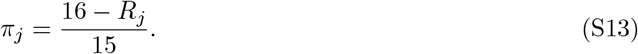

**BW-Podium:** The reward favours the three highest-ranked agents in the overall ranking:

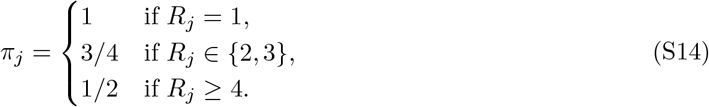

For all payment rules except `NC-Equal` and `NC-Score`, agents with identical scores receive the average of the rewards associated with the ranks they jointly occupy.

#### C.3 Convergence conditions and parameters of the genetic algorithm

The final stationary state results from the interplay of three mechanisms: selection, mutation, and genetic drift. Three conditions must be satisfied to ensure that the final state reflects the selective pressure of the payment rule.

The first condition is that the number of generations *N*_gen_ must be sufficiently large to allow the population to reach the final stable state. This convergence time must be several times larger than the number of generations required for the complete renewal of the entire population. At each generation, *N*_death_ agents die, of which a fraction *µ* are replaced by mutants. The mean number of mutants introduced per generation is *µN*_death_. The number of generations needed to renew the entire population through mutation is the inverse of the fraction of mutants introduced per generation:

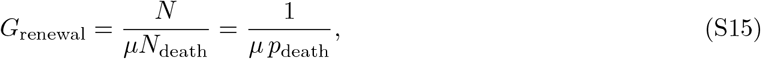

where *p*_death_ is the fraction of deaths per generation. The condition of convergence is thus *N*_gen_ ≫ *G*_renewal_. Between 5 and 10 times larger is typically enough.

The second and third conditions are that the fraction of mutants introduced per generation must be sufficiently large to prevent the fixation of a single profile by genetic drift, and sufficiently small to prevent driving the population towards the mutation spectrum. These two conditions are opposite on *µ*, but both must hold simultaneously.

The chosen parameters *N* = 100, *µ* = 0.1, *p*_death_ = 0.05, and *N*_gen_ = 1000 satisfy all three conditions simultaneously. The convergence condition gives *N*_gen_*/G*_renewal_ = 5, which is sufficient to reach the stationary state (see section D(f)). The fraction of mutants per generation is *µ p*_death_ = 0.005, small enough so that mutants do not dominate the population and, at the same time, the total number of mutants in the population is large enough to prevent fixation by genetic drift [46]. We verified that reducing *p*_death_ does not change the final stationary state, and that the results do not depend on the fraction *p*_parents_ = 0.4 of the population eligible as parents, provided it lies in the range 0.25–0.5. The stationary fractions are averaged over the last *N*_av_ = 50 generations of each run and over at least *N*_samp_ = 50 independent replicates to reduce Monte Carlo uncertainty.

The parameter *ε* = 0.1 controls the weight of the most recent games in the update of the fitness. This value ensures that the fitness reflects approximately the last ten games, while the contribution of games played before the current generation is negligible: 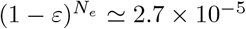 for *N*_*e*_ = 1000.

##### Computational efficiency and memory and disk usage

To reduce computational costs, we first generate a library of precomputed game outcomes by simulating 100 000 games for each of the 21 possible group compositions (e.g. CCCND, CCNND, NNNDD), storing the individual scores obtained in each game. During the genetic algorithm runs, the scores of agents in a given group are obtained by randomly sampling one of the 100 000 precomputed games corresponding to that group’s composition. This approach avoids repeatedly simulating the same types of interactions and substantially reduces computation time. Combined with just-in-time (JIT) compilation using Numba in Python, it allows approximately 7 × 10^10^ games to be sampled and processed within a few hours on a standard desktop computer.

Memory and disk requirements are further reduced by efficiently aggregating and storing results averaged over *N*_samp_ independent runs. This procedure preserves all the information needed to reproduce the figures without having to rerun the genetic algorithm.

The typical parameters used in the genetic algorithm are summarized in table S1.

**Table S1:** Parameters of the genetic algorithm.

| Parameter | Value | Description |
|---|---|---|
| *N* | 100 | Population size |
| *μ* | 0.1 | Mutation probability per individual and generation |
| *p*<sub>death</sub> | 0.05 | Fraction of individuals replaced per generation |
| *N*<sub>parents</sub> | 40% | Fraction of individuals eligible as parents |
| *N*<sub>gen</sub> | 1000 | Number of generations |
| *N*<sub>*e*</sub> | 1000 | Encounters per generation |
| *N*<sub>seeds</sub> | 50 | Independent replicates |

### D Supplementary genetic algorithm analyses

#### D.1 Genetic algorithm dynamics under a biased mutation spectrum

Figure S2 extends the analysis presented in figure 4 by introducing a biased mutation spectrum based on the behavioural frequencies measured experimentally under the `NC-Equal` payment rule, 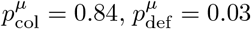. The relative ordering of the seven payment rules obtained in the unbiased case is preserved. This figure shows that in the absence of selective pressure the population converges to the mutation spectrum, confirming that the stationary states observed under the other payment rules are driven by selection and not by the mutation bias.

#### D.2 Parameter sweep: zero-mutation limit and robustness of rule ordering

We systematically varied the three parameters governing the genetic algorithm, namely the mutation rate *µ*, the fitness memory parameter *ε*, and the population renewal rate *p*_death_. The sweep over *µ* reveals the zero-mutation limit, in which the population tends towards fixation of a single strategy, depending on the payment rule. The sweeps over *µ, ε*, and *p*_death_ further show that the ordering of the seven payment rules by their capacity to sustain collaboration is preserved across the parameter ranges explored.

##### D.2.1 Zero-mutation limit

The sweep over *µ* reveals the asymptotic behaviour of the genetic algorithm as the mutation rate approaches zero. As *µ* → 0, the population tends towards fixation of a single behavioural strategy, whose identity depends on the fitness differences generated by the payment rule. `BG` favours fixation of collaborators. The four payment rules involving within-group competition, `WG`, `BW-Step`, `BW-Linear`, and `BW-Podium`, favour fixation of defectors. Under `NC-Equal`, different strategies can become fixed depending on stochastic fluctuations, whereas `NC-Score` favours fixation of neutrals.

The dependence of stationary strategy frequencies on *µ* is gradual over the investigated parameter range, without apparent discontinuities or changes in the relative ordering of the payment rules. Similar qualitative trends are observed for both initial population compositions (figure S3), indicating that the stationary strategy distributions are largely insensitive to the initial frequencies of behavioural strategies.

##### D.2.2 Robustness of payment-rule ordering

The relative ordering of the seven payment rules in their ability to promote and sustain cooperation remains largely unchanged across the explored parameter ranges. `BG` consistently produces the highest stationary fraction of collaborators, approaching 100% at low mutation rates and remaining above 65% at *µ* = 0.35. Conversely, `WG` strongly favours defection, with defectors dominating the stationary population. The three mixed-competition rules, `BW-Step`, `BW-Linear`, and `BW-Podium` produce intermediate levels of cooperation, generally in decreasing order. Under `NC-Equal`, stationary strategy frequencies remain close to the uniform mutation spectrum (1*/*3, 1*/*3, 1*/*3). Finally, neutrals dominate under `NC-Score`.

Varying *ε* and *p*_death_ further confirms the robustness of these results. The relative ordering of the payment rules is preserved throughout the parameter ranges explored, with no apparent qualitative changes in the stationary behavioural compositions. Similar stationary compositions are obtained from populations initially containing equal proportions of the three strategies and from populations composed entirely of collaborators. This convergence towards comparable outcomes suggests that the observed stationary regimes are largely independent of the initial population composition, rather than being artefacts of a particular initial condition.

#### D.3 Robustness of stationary strategy distributions

We assess the robustness of the stationary strategy distributions identified by the genetic algorithm. First, we examine whether similar stationary distributions are obtained from a broad range of initial population compositions. Second, we test their resilience to large perturbations (invasions) introduced after convergence.

##### D.3.1 Independence of initial conditions

To assess the independence of the GA outcomes on initial population composition, the GA is run under each payment rule using different initial conditions. These include homogeneous populations consisting entirely of collaborators, neutrals or defectors, mixtures of two strategies, and random mixtures of all three strategies. Across these configurations, the genetic algorithm converges towards similar stationary behavioural compositions for each payment rule. Figure S5 illustrates this convergence for three representative initial conditions.

**Figure S2:**
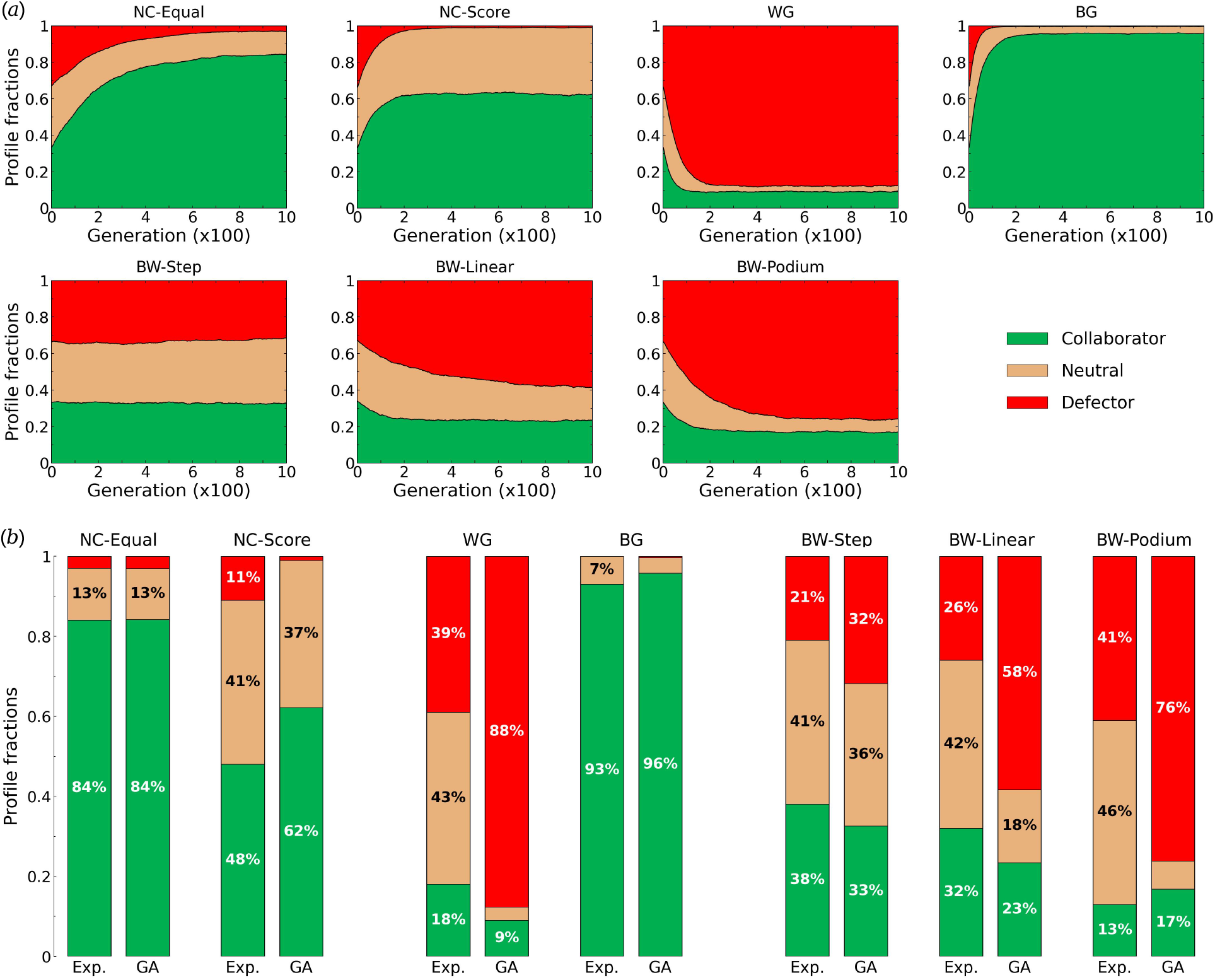
Genetic algorithm dynamics and comparison of stationary strategy distributions with experimental behavioural profiles under the seven payment rules, using a biased mutation spectrum. (*a*) Changes in strategy frequencies over successive generations of the genetic algorithm, starting from random initial strategy compositions. (*b*) Comparison of the asymptotic fractions of collaborators, neutrals and defectors obtained with the genetic algorithm (GA) with the corresponding fractions measured experimentally (Exp.) for each of the seven payment rules. Green areas show the fraction of collaborators *C*_*r*_, red areas show 1 ™ *D*_*r*_, where *D*_*r*_ is the fraction of defectors, brown area between both shows the fraction of neutrals 1 ™ *C*_*r*_ ™ *D*_*r*_. Parameter values of the GA: *N* = 100, *µ* = 0.1, *N*_gen_ = 1000, *ε* = 0.1, *p*_death_ = 0.05, *p*_parents_ = 0.40. Individuals are replaced using stochastic universal sampling (SUS). Mutations followed the experimentally derived distribution 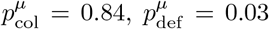, corresponding to the behavioural frequencies observed under `NC-Equal`. Average of *N*_samp_ = 500 replicas, final state averaged over the last *N*_av_ = 100 generations. See figure 4 in the main text for unbiased mutations.

**Figure S3:**
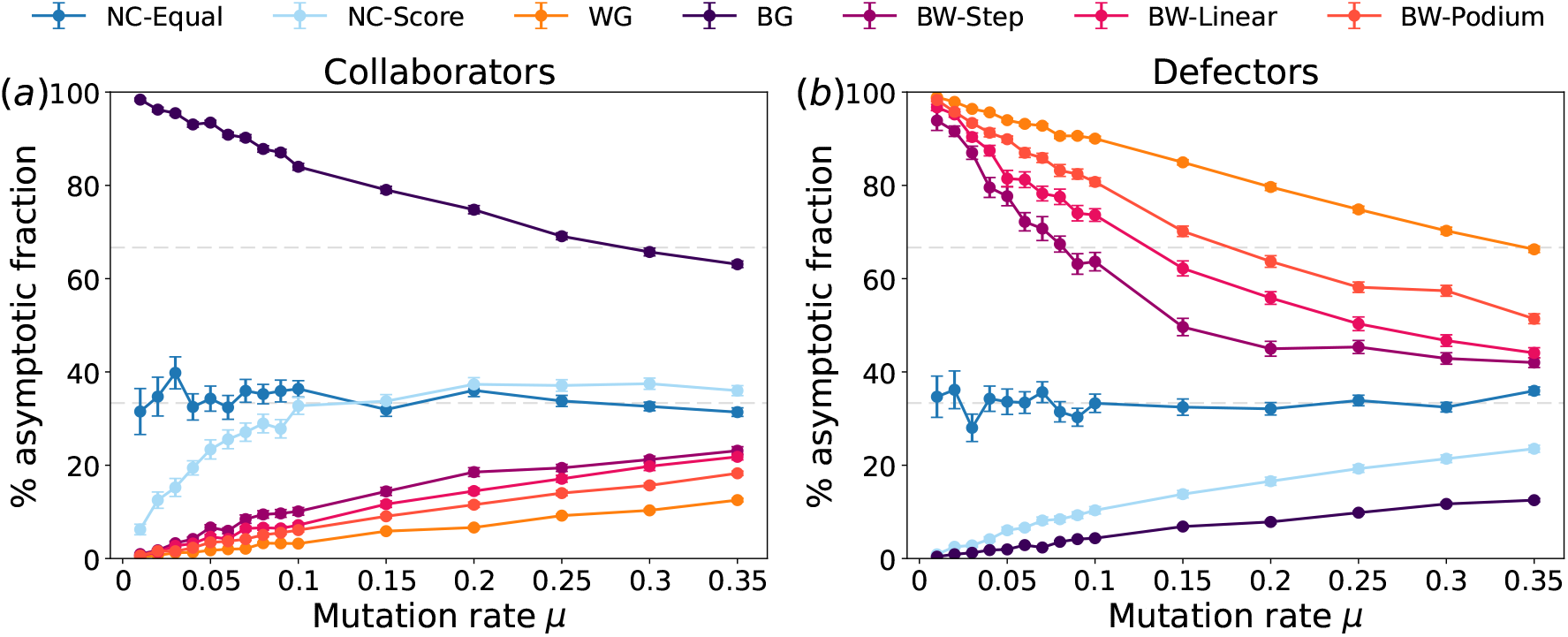
Effect of mutation rate on stationary behavioural compositions under the seven payment rules. Stationary fractions of (a) collaborators and (b) defectors as a function of the mutation rate *µ*, for each of the seven payment rules. Each point represents the mean of 50 independent runs and error bars indicate the standard error. Horizontal dashed lines indicate reference frequencies of 1/3 and 2/3. The populations initially contain equal proportions of collaborators, neutrals and defectors (1*/*3, 1*/*3, 1*/*3). Parameter values of the GA: *N* = 100, *ε* = 0.1, *p*_death_ = 0.05, *p*_parents_ = 0.40. The number of generations is *N*_gen_ = 5000 for *µ* ≤ 0.05 and *N*_gen_ = 1000 for *µ >* 0.05. Individuals are replaced using stochastic universal sampling (SUS). Mutations followed the unbiased spectrum 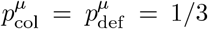. Average of *N*_samp_ = 50 replicas, final state averaged over the last *N*_av_ = 50 generations.

**Figure S4:**
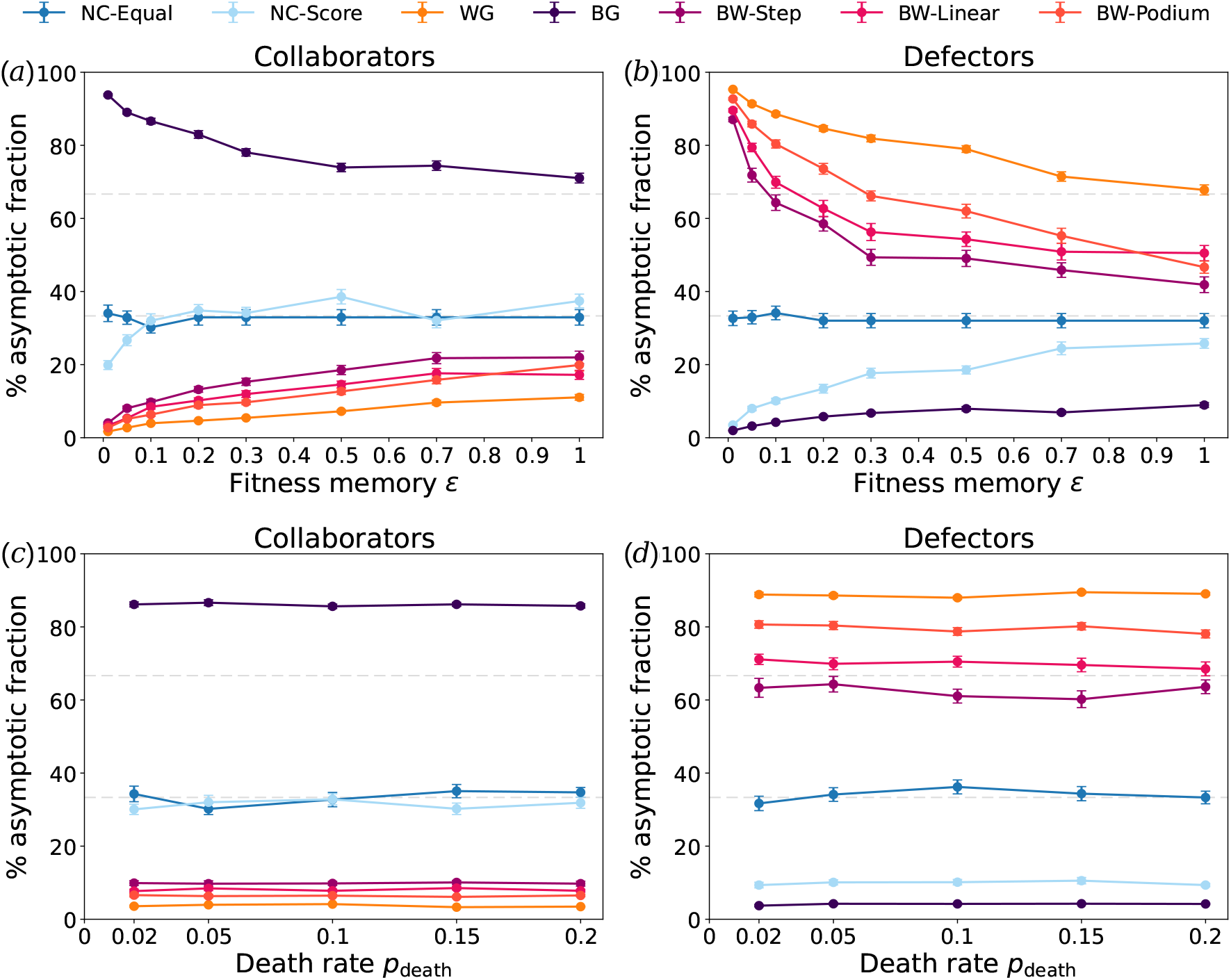
Robustness of stationary behavioural compositions to changes in fitness memory and population renewal rate. Stationary fractions of (*a,c*) collaborators and (*b,d*) defectors under the seven payment rules as functions of (*a,b*) the fitness memory parameter *ε* and (*c,d*) the death rate *p*_death_, for each of the seven rules. Each point represents the mean of 50 independent runs with error bars indicating the standard error of the mean. Horizontal dashed lines indicate reference frequencies of 1*/*3 and 2*/*3. Parameter values of the GA: (*a,b*) *p*_death_ = 0.05, *N*_gen_ = 8000 for *ε* = 0.01, *N*_gen_ = 5000 for *ε* = 0.05, *N*_gen_ = 3000 for *ε* = 0.1, *N*_gen_ = 2000 for *ε >* 0.1. (*c,d*) *ε* = 0.1, *N*_gen_ = 5000 for *p*_death_ = 0.02, *N*_gen_ = 3000 for *p*_death_ = 0.05, *N*_gen_ = 2000 for *p*_death_ *>* 0.05. Common values: *N* = 100, *µ* = 0.1, *p*_parents_ = 0.40. Individuals are replaced using stochastic universal sampling (SUS). Mutations followed the unbiased spectrum 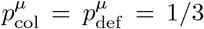. Average of *N*_samp_ = 50 replicas, final state averaged over the last *N*_av_ = 50 generations.

**Figure S5:**
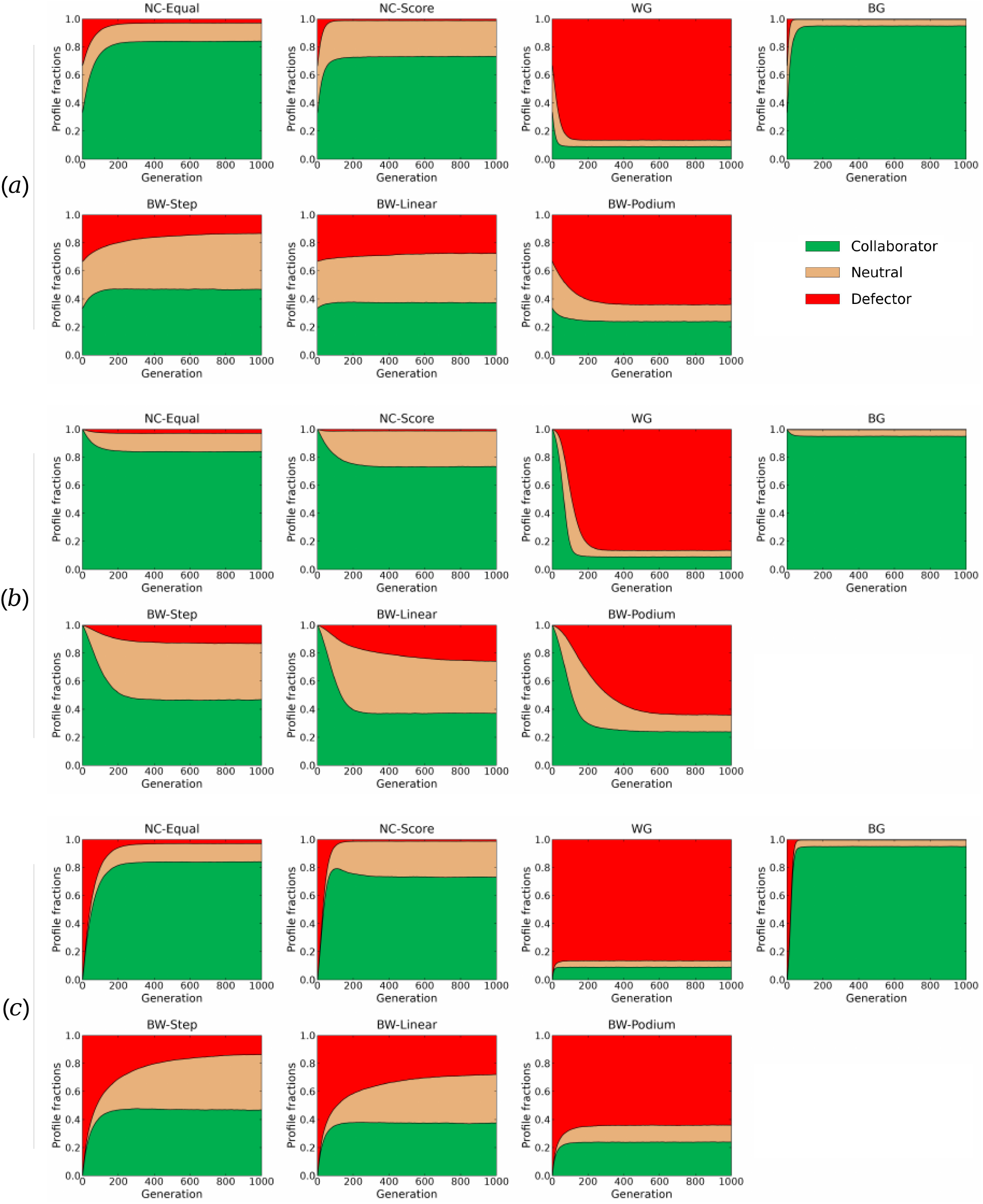
Convergence towards stationary behavioural compositions from different initial conditions. Changes in the frequencies of collaborators, neutrals and defectors over successive generations of the genetic algorithm under the seven payment rules, starting from (*a*) equal initial proportions of the three strategies, (*b*) a population composed entirely of collaborators, and (*c*) a population composed entirely of defectors. Agents are randomly assigned to groups before each game. Parameter values of the GA: *N* = 100, *µ* = 0.35, *ε* = 0.1, *p*_death_ = 0.05, *p*_parents_ = 0.4. Mutations followed the experimentally derived distribution 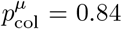, 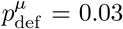, corresponding to the behavioural frequencies observed under `NC-Equal`. Each payment-rule condition is averaged over *N*_samp_ = 5000 independent runs, corresponding to ×10 five-player games. Standard errors of the final strategy frequencies are negligible at the scale of the plots and are therefore not shown. For each payment rule, the three initial conditions converge similar stationary plateaux within *N*_gen_ = 1000 generations.

##### D.3.2 Robustness to strategy invasions

To assess the robustness of the stationary behavioural compositions identified by the genetic algorithm, we perform invasion experiments in which established populations are subject to large perturbations. After an initial evolutionary period of 2000 generations, 60% of the population is replaced by agents adopting an invading behavioural strategy. The genetic algorithm is then run for a further 2000 generations to determine whether the population recovers its previous stationary composition or converges towards a different state. In all cases, the populations return towards their characteristic stationary compositions under the corresponding payment rules, indicating that these compositions are robust to substantial changes in strategy frequencies. Figure S6 shows three representative scenarios: collaborators invading a defector population, defectors invading a collaborator population, and neutrals invading a collaborator population.

---

**Figure S6:**
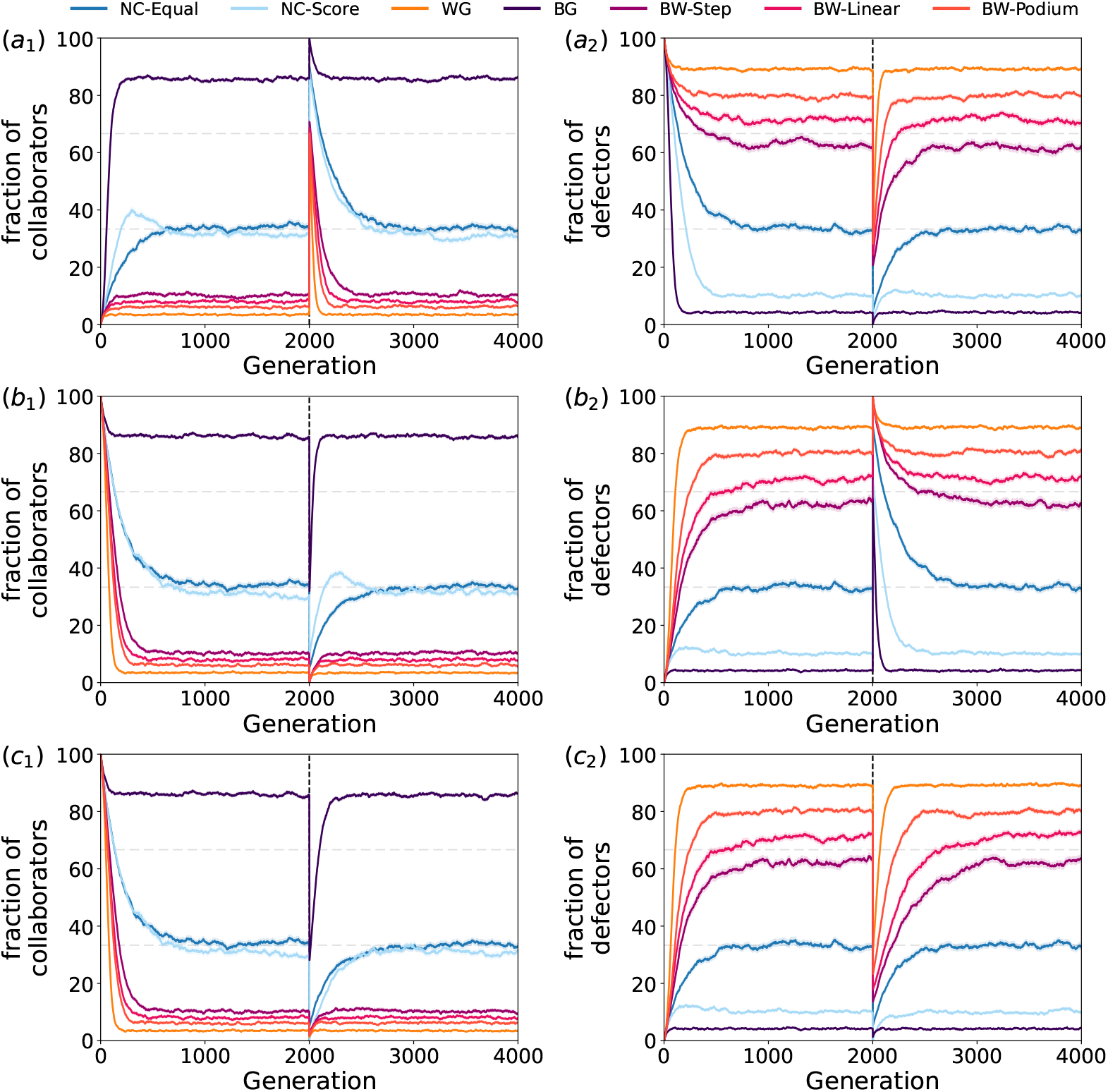
Robustness of stationary behavioural compositions to large strategy invasions. (*a*) Collaborators invading a population initially composed of defectors, (*b*) defectors invading a population initially composed of collaborators, and (*c*) neutrals invading a population initially composed of collaborators. In each scenario, the invasion occurred after 2000 generations (vertical dashed line), and is continued for another 2000 generations. (*a*_1_, *a*_2_) Following the introduction of collaborators, populations returned towards their pre-invasion stationary compositions under all seven payment rules. (*b*_1_, *b*_2_) Similarly, populations recovered their characteristic stationary compositions following the introduction of defectors, including under the strongly cooperation-promoting `BG` rule. (*c*_1_, *c*_2_) The introduction of neutrals also produced only transient changes, with populations subsequently returning towards their characteristic stationary compositions. These results indicate that the stationary behavioural compositions are resilient to substantial perturbations in population composition. Parameters of the genetic algorithm are *N* = 100, *µ* = 0.1, *ε* = 0.1, *p*_death_ = 0.05. Agents are selected for replacement using stochastic universal sampling (SUS), with a uniform mutation spectrum. Each phase lasts *N*_gen_ = 2000 generations, and the results are averaged over 200 independent runs. At the time of invasion (*G* = 2000), invading agents are assigned the maximum fitness of the resident population.

